# Online analysis of microendoscopic 1-photon calcium imaging data streams

**DOI:** 10.1101/2020.01.31.929141

**Authors:** Johannes Friedrich, Andrea Giovannucci, Eftychios A. Pnevmatikakis

## Abstract

In-vivo calcium imaging through microendoscopic lenses enables imaging of neuronal populations deep within the brains of freely moving animals. Previously, a constrained matrix factorization approach (CNMF-E) has been suggested to extract single-neuronal activity from microendoscopic data. However, this approach relies on offline batch processing of the entire video data and is demanding both in terms of computing and memory requirements. These drawbacks prevent its applicability to the analysis of large datasets and closed-loop experimental settings. Here we address both issues by introducing two different online algorithms for extracting neuronal activity from streaming microendoscopic data. Our first algorithm presents an online adaptation of the CNMF-E algorithm, which dramatically reduces its memory and computation requirements. Our second algorithm proposes a convolution-based background model for microendoscopic data that enables even faster (real time) processing on GPU hardware. Our approach is modular and can be combined with existing online motion artifact correction and activity deconvolution methods to provide a highly scalable pipeline for microendoscopic data analysis. We apply our algorithms on two previously published typical experimental datasets and show that they yield similar high-quality results as the popular offline approach, but outperform it with regard to computing time and memory requirements.

**Author summary:** Calcium imaging methods enable researchers to measure the activity of genetically-targeted large-scale neuronal subpopulations. Whereas previous methods required the specimen to be stable, e.g. anesthetized or head-fixed, new brain imaging techniques using microendoscopic lenses and miniaturized microscopes have enabled deep brain imaging in freely moving mice.

However, the very large background fluctuations, the inevitable movements and distortions of imaging field, and the extensive spatial overlaps of fluorescent signals complicate the goal of efficiently extracting accurate estimates of neural activity from the observed video data. Further, current activity extraction methods are computationally expensive due to the complex background model and are typically applied to imaging data after the experiment is complete. Moreover, in some scenarios it is necessary to perform experiments in real-time and closed-loop – analyzing data on-the-fly to guide the next experimental steps or to control feedback –, and this calls for new methods for accurate real-time processing. Here we address both issues by adapting a popular extraction method to operate online and extend it to utilize GPU hardware that enables real time processing. Our algorithms yield similar high-quality results as the original offline approach, but outperform it with regard to computing time and memory requirements. Our results enable faster and scalable analysis, and open the door to new closed-loop experiments in deep brain areas and on freely-moving preparations.

## Introduction

In vivo calcium imaging of activities from large neural populations at single cell resolution has become a widely used technique among experimental neuroscientists. Recent advances in optical imaging technology using a 1-photon-based miniscope and a microendoscopic lens have enabled in vivo calcium imaging studies of neural activities in freely behaving animals [1–3]. However, this data typically displays large, blurry background fluctuations due to fluorescence contributions from neurons outside the focal plane, arising from the large integration volume of one photon microscopy. To obtain a robust approach for extracting single-neuronal signals from microendoscopic data the constrained nonnegative matrix factorization (CNMF, [4]) approach has been extended to leverage a more accurate and flexible spatio-temporal background model able to capture the properties of the strong background signal (CNMF-E, [5]). This prevalent algorithm (see [6] for an alternative proposal) has been widely used to study neural circuits in cortical and subcortical brain areas, e.g. prefrontal cortex (PFC) and hippocampus [5], as well as previously inaccessible deep brain areas, such as striatum [7], amygdala [8], substantia nigra pars compacta (SNc) [9], nucleus accumbens [10], dorsolateral septum [11], parabrachial nucleus [12], and other brain regions.

A concomitant feature of the refined background model in CNMF-E is its high computational and memory cost. Although the data can be processed by splitting and processing the FOV in smaller patches to exploit a time/memory tradeoff [13], this strategy requires significant time resources, does not scale to longer recordings, and introduces border effects among patches when estimating the background. Further, CNMF-E is applied to imaging data after the experiment is complete. However, in many cases we would prefer to run closed-loop experiments – analyzing data on-the-fly to guide the next experimental steps or to control feedback [14–16] – and this requires new methods for accurate real-time processing.

Online (and real time) analysis of calcium imaging data has been proposed with the OnACID algorithm [17]. The algorithm combines the online NMF algorithm of [18], the CNMF source extraction algorithm of [4], and the near-online deconvolution algorithm of [19], to provide an automated pipeline that can discover and track the activity of hundreds of cells in real time, albeit only for 2-photon or light-sheet imaging data.

In this paper, we present two algorithms for the online analysis of microendoscopic 1-photon calcium imaging data streams. Our first algorithm (OnACID-E), extends [17] by incorporating the background model and neuron detection method of CNMF-E [5] and adapting them to an online setup. Our second approach proposes a lower dimensional background model by introducing parameter sharing through a convolutional structure and combines it with the online 2-photon processing of [17]. In either approach, every frame is processed in four sequential steps: i) The frame is registered against the previous background-corrected denoised frame to correct for motion artifacts. ii) The fluorescence activity of the already detected sources is tracked. iii) Newly appearing neurons and processes are detected and incorporated to the set of existing sources. iv) The fluorescence trace of each source is denoised and deconvolved to provide an estimate of the underlying spiking activity.

Our resulting framework is highly scalable with minimal memory requirements, as it processes the data in streaming mode (one frame at a time), while keeping in memory a set of low dimensional sufficient statistics and a small minibatch of the most recent data frames. Moreover, it results in faster processing that can reach real time speeds for common experimental scenarios when utilizing GPU hardware. We apply our framework to typical mouse *in vivo* microendoscopic 1p datasets; our algorithm can find and track hundreds of neurons faster than real-time, and outperforms the CNMF-E algorithm of [5] with regard to computing time and memory requirements while maintaining the same high quality of the results. We also provide a Python implementation of our methods as part of the CaImAn package [13].

## Methods

This section is organized as follows. The first subsection briefly reviews the modeling assumptions of CNMF-E for microendoscope data. In the second subsection, we derive an online method to fit this model, thus enabling the processing of 1-photon endoscopic data streams (OnACID-E). In the third subsection, we modify the background modeling assumptions to introduce a convolutional structure and describe how to utilize this to derive an alternative fast online algorithm. Finally, we describe how motion correction, which is typically done as preprocessing step, can not only be performed online as well, but even profit from stream processing.

### CNMF for microendoscopic data (CNMF-E)

The recorded video data can be represented by a matrix 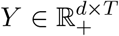, where *d* is the number of imaged pixels and *T* is the number of frames observed. Following [4], we model *Y* as

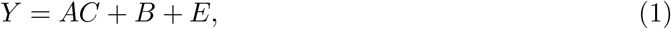

where 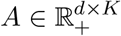 is a spatial matrix that encodes the location and shape of each neuron (spatial footrpint), 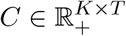 is a temporal matrix that characterizes the fluorescence of each neuron over time, matrix *B* represents background fluctuations and *E* is additive Gaussian noise with mean zero and diagonal covariance.

The CNMF framework of [4] incorporates further constraints beyond non-negativity. Each spatial footprint **a**_*i*_ is constrained to be spatially localized and hence sparse. Similarly, the temporal components **c**_*i*_ are highly structured, as they represent the cells’ fluorescence responses to typically sparse, nonnegative trains of action potentials. Following [19, 20], we model the calcium dynamics of each neuron **c**_*i*_ with a stable autoregressive process of order *p*,

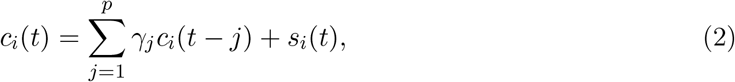

where *s*_*i*_(*t*) ≥ 0 is the number of spikes that neuron *i* fired at the *t*-th frame, and *γ*_*j*_, *j* = 1, …, *p* correspond to the discrete time constants of the dynamics that depend on the kinematic properties of the used indicator.

For the case of microendoscopic data the background is modeled as [5]

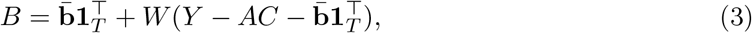

where **1**_*T*_ denotes a vector of *T* ones, 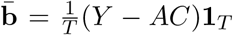 models constant baselines and the second term fluctuating activity. *W* is an appropriate sparse weight matrix, where *W*_*ij*_ models the influence of the neuropil signal of pixel *j* to the neuropil signal at pixel *i*. It is constrained to *W*_*ij*_ = 0 if dist(**x**_*i*_, **x**_*j*_) ∉ [*l, l* + 1[, thus we model the background at one pixel as a linear combination of the background fluorescence in pixels which are chosen to be on a ring with radius *l*. Typically, *l* is chosen to be ~1.5× the radius of an average neuron, to exclude contributions that might be affected from the activity of an underlying neuron.

### Fitting the CNMF-E model

We first recap the offline approach for fitting the CNMF-E model [5], and then show how it can be adapted to an online setup.

### Offline

The estimation of all model variables can be formulated as a single optimization problem

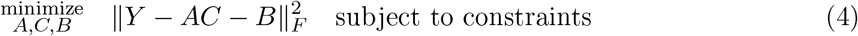

The CNMF-E algorithm of [5] divides the nonconvex problem (4) into three simpler subproblems that are solved iteratively: Estimating *A* given estimates *C* and *B*, estimating *C* given *A* and *B*, and estimating *B* given *A* and *C*.

*A* and *C* are estimated using a modified version of “fast hierarchical alternating least squares” [21] that includes sparsity and localization constraints [22]. The update of *A* consists of block-coordinate decent steps iterating over neurons *i*,

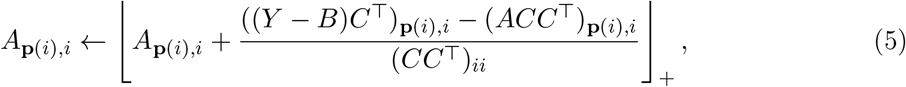

where **p**(*i*) specifies the pixel indices where *A*_:*,i*_ can take non-zero values, i.e. where neuron *i* is located. For computational efficiency the sufficient statistics *L* = (*Y − B*)*C*^T^ and *M* = *CC*^T^ are computed only once initially and cached.

Similarly, the block-coordinate decent steps for updating *C* are

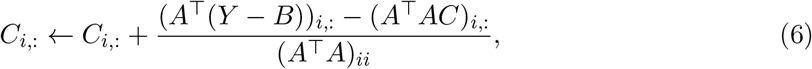

with sufficient statistics *A*^T^(*Y − B*) and *A*^T^*A* computed only once initially. *C* should not merely be constrained to non-negative values but follow the dynamics of the calcium indicator, thus to further denoise and deconvolve the neural activity from the dynamics of the indicator the OASIS algorithm [19] is used. OASIS solves a modified LASSO problem

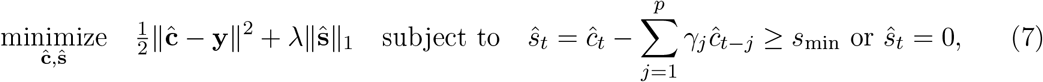

where **y** denotes a noisy neural calcium trace obtained as result of Eq (6). The *l*_1_ penalty on 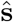 or the minimal spike size *s*_min_ can be used to enforce sparsity of the neural activity.

The spatiotemporal background is estimated from the linear regression problem

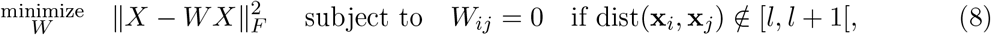

where 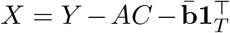 and 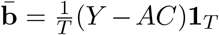. The solution is given by the normal equations for each pixel *i*,

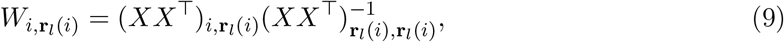

where **r**_*l*_(*i*) = {*j*|dist(**x**_*i*_, **x**_*j*_) ∈ [*l, l* + 1[} specifies the pixel indices where *W*_*i,*:_ can take non-zero values. Given the optimized *W*, the whole background signal is 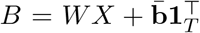. More information can be found in [5].

### Online

The offline framework presented above can be adapted to a data streaming setup by appropriately modifying the online NMF algorithm of [18], and the online algorithm for analyzing 2-photon calcium imaging data [17]. Using Eq (1), the observed fluorescence at time *t* can be written as

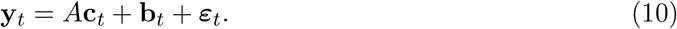

The (non-deconvolved) activity of all neurons at time *t*, **c**_*t*_, is obtained by iteratively evaluating Eq (6) given raw frame data **y**_*t*_, spatial footprints *A*, and background parameters *W,* 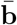.The activity is further denoised and deconvolved by running OASIS [19], which is not only a very fast algorithm, but crucially progresses through each time series sequentially from beginning to end and is thus directly applicable to stream processing. The background term in Eq (6) evaluates to 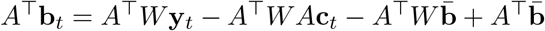 and for computational efficiency the terms *A*^T^*W*, *A*^T^*W A* and 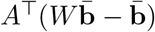 are maintained in memory and updated incrementally, cf. Alg S1 in Supplementary Material. Warm starts are exploited by initializing **c**_*t*_ with the value at the previous frame **c**_*t*−1_, since the calcium traces *C* are continuous and typically change slowly. Moreover, the temporal traces of components that do not spatially overlap with each other can be updated simultaneously in vector form; we use a simple greedy scheme to partition the components into spatially non-overlapping groups [17].

The spatial footprints *A* are obtained by iteratively evaluating Eq (5) and can be estimated efficiently as in [18] by only keeping in memory the sufficient statistics

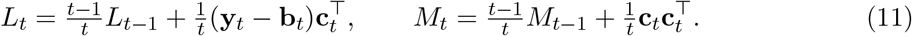

Since neurons’ shapes are not expected to change at a fast timescale, updating *A* is actually not required at every timepoint; in practice we update every 200 time steps, again warm started at the value from the previous iteration, cf. Alg 1. Additionally, the sufficient statistics *L*_*t*_, *M*_*t*_ are only needed for updating the estimates of *A* so they can be updated only when required. Further, Eq (5) accesses only elements **p**(*i*) in column *i* of *L*, hence only those entries of *L* need to be updated, cf. Algs S2 and S3 in Supplementary Material.

To update the background components *W,* 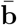, we keep track of the constant baselines 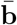 and the sufficient statistics χ = *XX*^*T*^ that is needed to compute *W* using Eq (9)

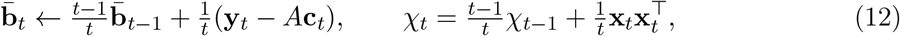

where 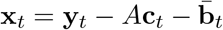. As is the case with the spatial footprints, updating the background is actually not required at every timepoint and in practice we update every 200 time steps, cf. Alg 1 and Alg S2 in Supplementary Material. Processing pixel *i* according to Eq (9) (see also Alg S4 in Supplementary Material) accesses only vector 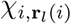 and sub-matrix 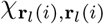. Some elements of *χ* are not part of any sub-matrix or vector for any *i* and thus are never accessed. In practice we therefore update and store only these vectors and sub-matrices for computational and memory efficiency. Because the background has no high spatial frequency components, it can be spatially decimated to further speed up processing [19] without compromising the quality of the results. E.g. downscaling by a factor of 2 reduces the number of pixels by a factor of 4 and the number of elements in *W* and *χ* by a factor of 16. Less and smaller least squares problems (Eq 9) need to be solved, which drastically reduces processing time and memory consumption.

Note that updating the background components and all the spatial footprints at a given frame results in a computational bottleneck for that specific frame. While on average, this effect is minimal (cf. Results section and Fig 4) a temporary slowdown can have an adverse effect on a real-time closed loop setup. This restriction can be lifted by holding the background model fixed and updating the spatial footprints in a distributed manner across all frames. As described, later, using a lower dimensional background model can achieve that and enable fast real time processing with balanced workload across all frames.

**Algorithm 1.**
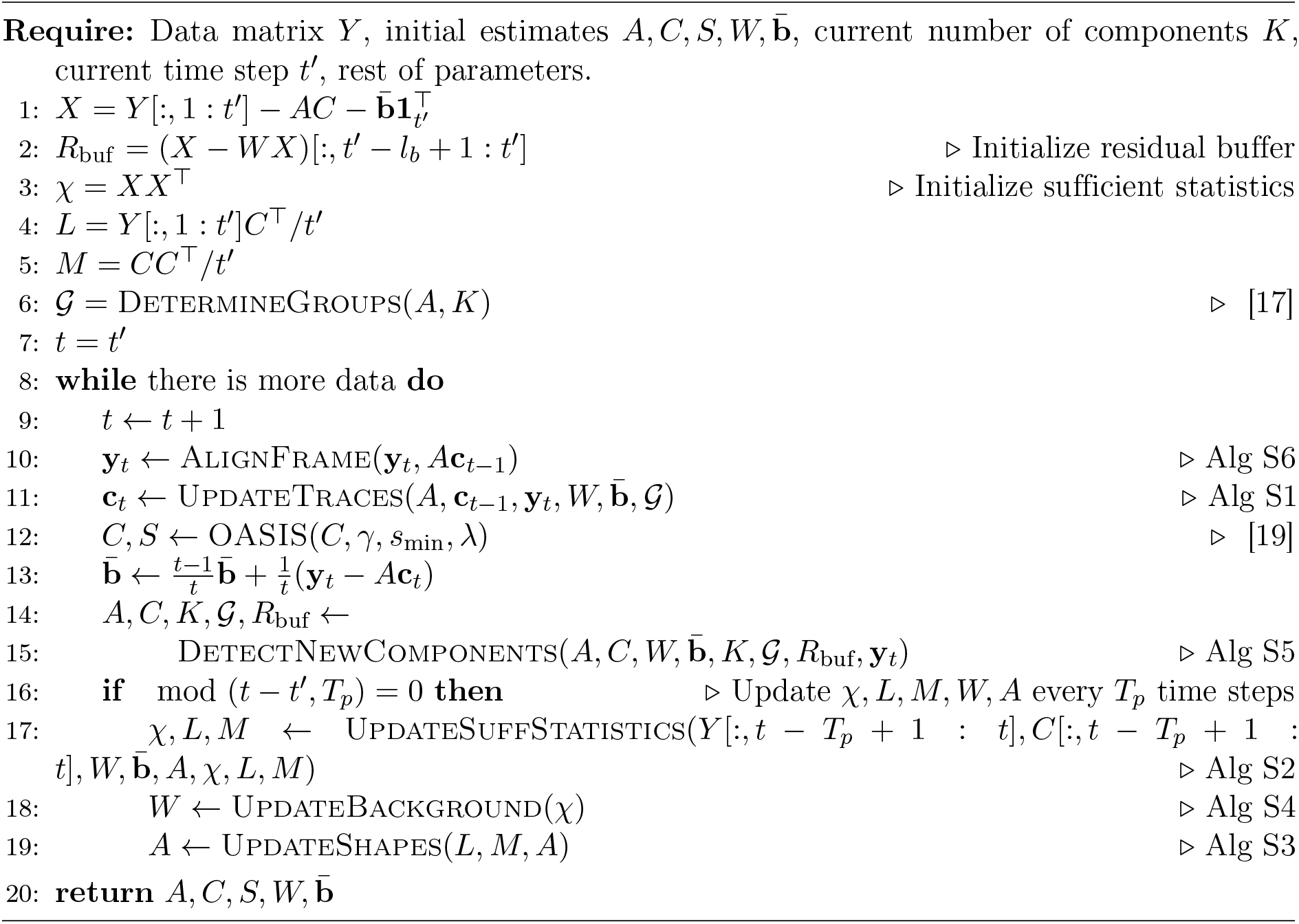
OnACID-E

**Fig 4.**
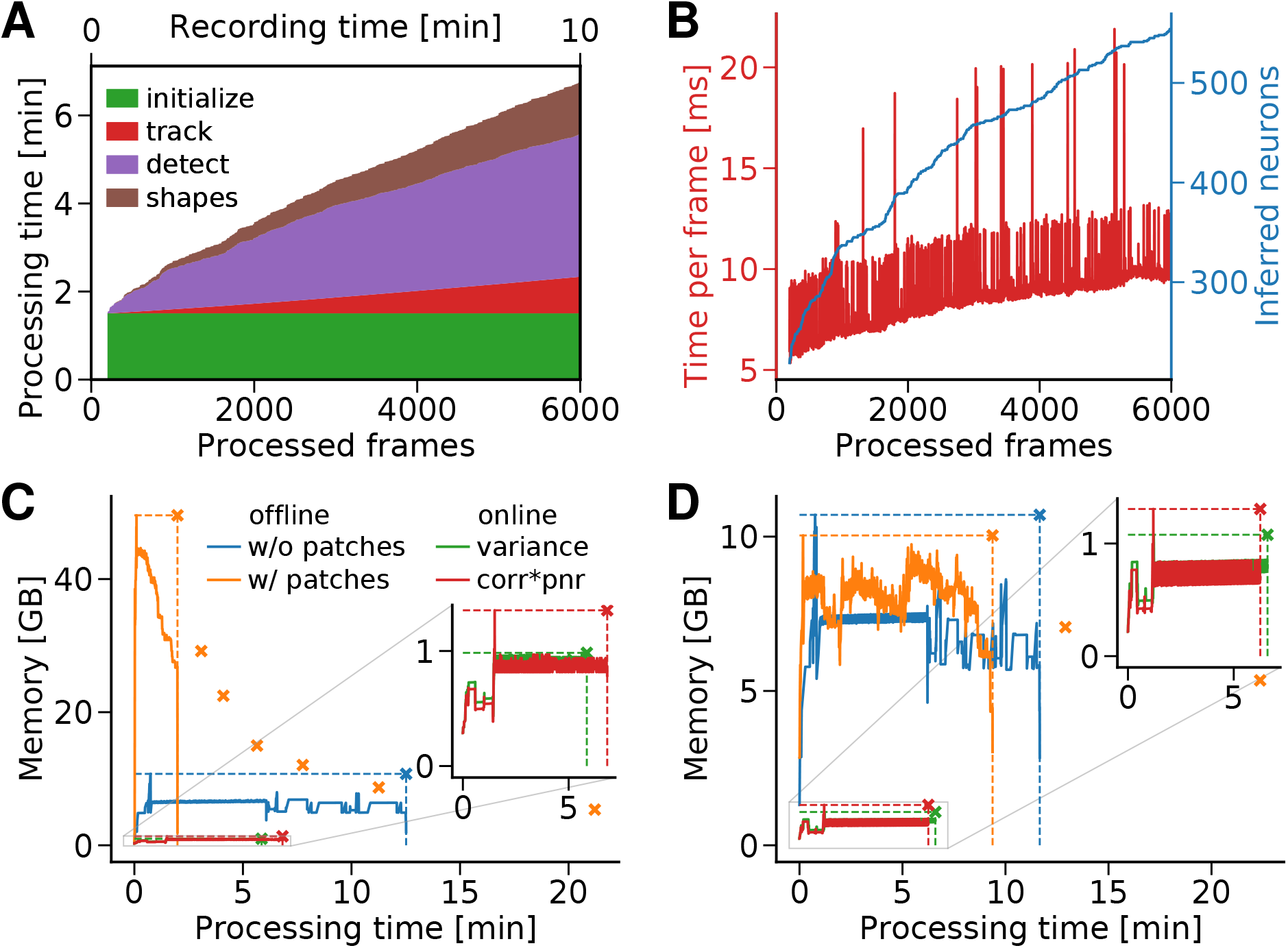
Computing resources of OnACID-E. **(A)** Cumulative processing time, separated by time for initialization (occurred only at the beginning), tracking existing activity, detecting new neurons, and updating spatial footprints as well as background. **(B)** Cost of tracking neurons’ activity. **(C)** Memory consumption of OnACID-E and CNMF-E. Markers and dashed lines indicate peak memory and overall processing time, whereas solid lines show memory as function of time. Offline processing using CNMF-E was performed with or without patches, online processing using OnACID-E with variance or corr☉pnr summary image, cf. Methods. The orange markers show peak memory and overall processing time when the number of parallel processes is varied (16, 8, 6, 4, 3, 2 and 1), illustrating the time-memory trade off when processing in patches (more processes can lead to faster processing at the expense of additional memory requirements). **(D)** Analogous results as in (C) when using a laptop. The dataset consisted of 6000 frames with a 256×256 FOV.

To initialize our algorithm we use the CNMF-E algorithm on a short initial batch of data of length *T*_*b*_, (e.g., *T*_*b*_ = 200). The sufficient statistics are initialized from the components that the offline algorithm finds according to Eqs (11, 12).

### Detecting new components

The approach explained above enables tracking the activity of a fixed number of sources, and will ignore neurons that become active later in the experiment. Following [17], we approach the problem by introducing a buffer that contains the last *l*_*b*_ instances of the residual signal **r**_*t*_ = **y**_*t*_ − *A***c**_*t*_ − **b**_*t*_, where *l*_*b*_ is a reasonably small number, e.g., *l*_*b*_ = 100. From this buffer we compute a summary image (as detailed later we actually update the summary image instead of computing it afresh) and then search for the local maxima of the image to determine new candidate neurons.

One option for the summary image **e** is to proceed along the lines of [4], i.e. to perform spatial smoothing with a Gaussian kernel with radius similar to the expected neuron radius, and then calculate the energy for each pixel *i*, 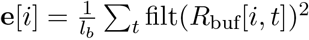, where filt() refers to the smoothing operation. Another option is to follow [5] and calculate the peak-to-noise ratio (PNR),

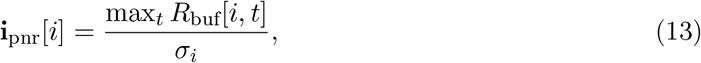

as well as the local cross-correlation image,

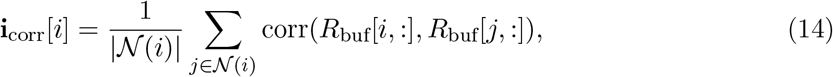

where 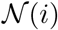 specifies the neighboring pixels of pixel *i* and the function corr() refers to Pearson correlation. Their pixel-wise product **e**= **i**_pnr_ ☉ **i**_corr_ is used as summary image. We use the latter throughout the Results section, if not explicitly stated otherwise. New candidate components **a**_new_, and **c**_new_ are estimated by performing a local rank-1 NMF of the residual matrix restricted to a fixed neighborhood around the point of maximal variance, or maximal product of PNR and cross-correlation, respectively.

To limit false positives, the candidate component is screened for quality. Similarly to [17], to prevent noise overfitting, the shape **a**_new_ must be significantly correlated (e.g., *r* ~ 0.5) to the residual buffer averaged over time and restricted to the spatial extent of **a**_new_. Moreover, if **a**_new_ significantly overlaps with any of the existing components, then its temporal component **c**_new_ must not be highly correlated with the corresponding temporal components; otherwise we reject it as a possible duplicate of an existing component. Once a new component is accepted, *A, C* are augmented with **a**_new_ and **c**_new_ respectively, the quantities *A*^T^*W*, *A*^T^*WA* and 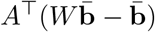 are updated via augmentation, and the sufficient statistics are updated as follows:

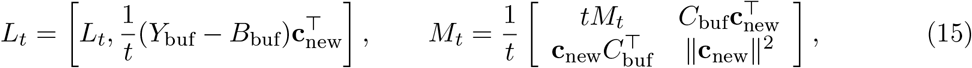

where 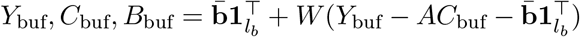 denote the matrices *Y, C, B*, restricted to the last *l*_*b*_ frames that the buffer stores. This process is repeated until no new components are accepted, at which point the next frame is read and processed.

### Updating the summary image

For computational efficiency we avoid repeated computations and perform incremental updates of the summary image instead of computing it afresh. If the variance image is used, it is updated according to 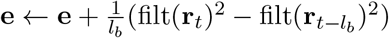 when the next frame is processed. When a new component with footprint **a** is added the residual changes at the component’s location and we update the variance image accordingly locally only for pixels *i* where the smoothed component is positive (filt(**a**)[*i*] > 0) according to 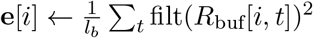

Next we consider the case that the product of cross-correlation image and PNR image is used as summary image. We keep track of the first and second order statistics

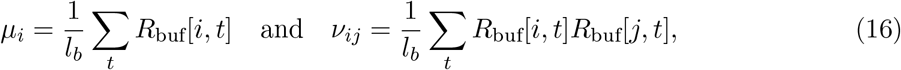

the latter only for pixels 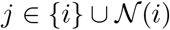. These statistics are updated according to

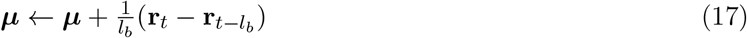

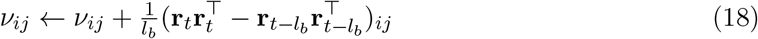

when the next frame is processed. The cross-correlation values are computed from these statistics as

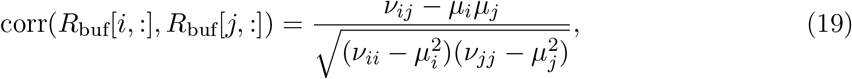

and the correlation image is obtained according to Eq (14). For computing the PNR image we use the noise level *σ*_*i*_ estimated on the small initial batch for the denominator in Eq (13) and keep track of the maximum image **i**_max_ ← max(**i**_max_, **r**_*t*_) for the nominator. When a new component with foot print **a** and time series 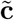 is added we set **i**_max_[*i*] to zeros if *a*_*i*_ > 0. The statistics for the cross-correlation are updated as

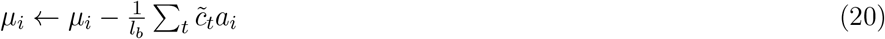

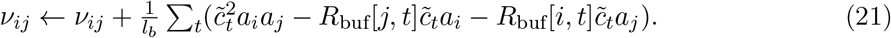

The whole online procedure of OnACID-E is described in Algorithm 1; Supplementary Material includes pseudocode description of the referenced routines.

### Background modeling using convolutional neural networks

The background model used in the CNMF-E algorithm (Eq (8)) assumes that the value of the background signal at a given point in space is given by a linear combination of the background values from the points in a ring centered around that pixel with width 1 and radius *l*, where *l* is larger than the radius of the typical neuron in the dataset by a small factor (e.g. 1.5) plus a pixel dependent scalar [5]. While powerful in practice, this model does not assume any dependence between the linear combination weights of all the different pixels, and results in a model with a very large number of parameters to be estimated. Ignoring pixels near the boundary, each row of the matrix *W* which represents the linear combination weights will have approximately [2*πl*] non-zero entries (where [·] denotes the integer part), giving a total number of *d*([2*πl*] + 1) parameters to be estimated. While this estimation can be done efficiently in parallel as discussed above, and the overall number of parameters can be reduced through spatial downsampling, we expect that the overall number of degrees of freedom in such a model is much lower. The reason is that the “ring” model aims to capture aspects of the point spread function which is largely invariant with respect to the location within the FOV.

To test this hypothesis we used a very simple convolutional neural network (CNN) with ring shaped kernels to capture the background structure. The intuition behind the convolution is straightforward: if all the rows of the *W* matrix had the same non-zero entries (but centered around different points) then the application of *W* would correspond to a simple spatial convolution with the common “ring” as the filter. In our case this is not sufficient and therefore we investigated parametrizing the background model with a slightly more complex model, which we refer to as “Ring-CNN”.

Let *f*_***θ***_ : ℝ^*d*^ → ℝ^*d*^ be a function that models the autoregressive nature of the background. In the CNMF-E case this simply corresponds to 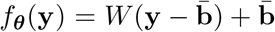. In the linear model we parametrize the function as

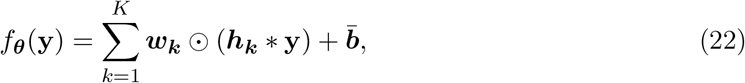

where ***b***, ***w***_*k*_ ∈ ℝ^*d*^, *k* = 1, …, *K*, and ☉,∗ refer to pointwise multiplication and spatial convolution, respectively (with slight abuse of notation we assume that **y** has been reshaped back to 2d image to perform the convolution and the result of the convolution is again vectorized). Finally, ***h***_***k***_, *k* = 1, …, *K* is a ring shaped convolutional kernel which takes non-zero values only at a specified annulus around its center. Note that this corresponds to parametrizing directly W as

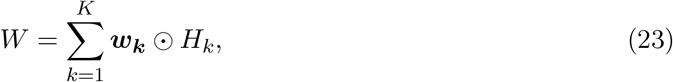

where *H*_*k*_ ∈ ℝ^*d*×*d*^ is the matrix induced by the convolutional kernel ***h***_***k***_. Intuitively this model corresponds to using a pixel dependent linear combination of *K* ring basis functions, and results in a total *K*(*d* + [2*πl*]) + *d* parameters to be estimated. Compared to the *d*([2*πl*] + 1) number of parameters for the CNMF-E model, this can result in a significant reduction when *K* < [2*πl*].

Note that decoupling the number of different “rings” from the total number of pixels, enables the consideration of wider “rings” that integrate over a larger area of the FOV and can potentially provide more accurate estimates, without a dramatic increase on the number of parameters to be learned. For example, a “ring” with inner radius *l* and width *w* would require approximately [*πw*(2*l* + *w* − 1)] parameters and the total number of parameters would be *K*([*πw*(2*l* + *w* − 1)] + *d*) as opposed to *d*([*πw*(2*l* + *w* − 1)] + 1) for the standard CNMF-E model.

### Unsupervised training on the raw data

To estimate the autoregressive background model in the CNMF-E algorithm, we want to operate on the data after the spatiotemporal activity of all detected neurons has been removed (Eq (8)). For the CNN model this would translate into the optimization problem

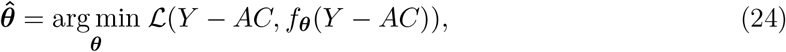

where 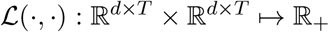 is an appropriate loss function (e.g. the Frobenius norm).

For the CNMF-E algorithm, operating on *Y − AC* is necessary because each “ring” has its own independent weights whose estimation can be biased from the activity of nearby neurons. In the CNN case however, the background model assumes a significant amount of weight sharing between the different “rings” which makes the estimation more robust to the underlying neural activity. Therefore we can estimate the background model by solving directly

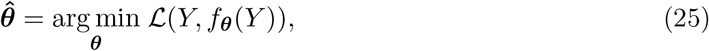

meaning that the solution of Eq (25) should satisfy 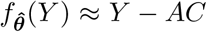. This approximation is based on the assumption that the activity of neurons is sparse so the product *AC* will be much smaller than *f*_***θ***_(*Y*) most of the time. To promote this we can use the *L*_1_ norm of the difference as the loss function 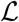. Furthermore, since *AC* is nonnegative we seek to under-approximate *Y* with the background *f*_***θ***_(*Y*). To encode that in the objective function we can consider a quantile loss function [23] that penalizes over-approximation more than under-approximation:

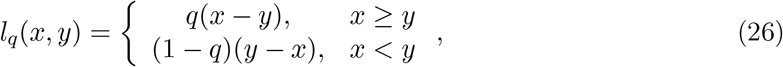

For some *q* ∈ (0, 1] and take 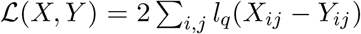. For example, for *q* = 0.5 Eq (26) corresponds to the *L*_1_ norm of the difference, and to promote the under-approximation property we use *q* < 0.5. Since the models are differentiable and the objective function is additive, Eq (25) can be optimized in an online mode using stochastic gradient descent.

### Online Processing

In practice, we found that by using rings of increased width (e.g. 5 pixels), training the model only during the initialization process on a small batch frames, leads to convergence due to the large amount of weight sharing that reduces the number of parameters. Once the model has been trained, it can be used to remove the background from the data (after motion correction). To reduce the effect of active neurons on the inferred background we can approximate the activity at time *t*, with the activity at time *t* − 1, and subtract that from the data frame prior to computing the background. In other words, we can use the approximation

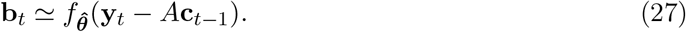

**Algorithm 2.**
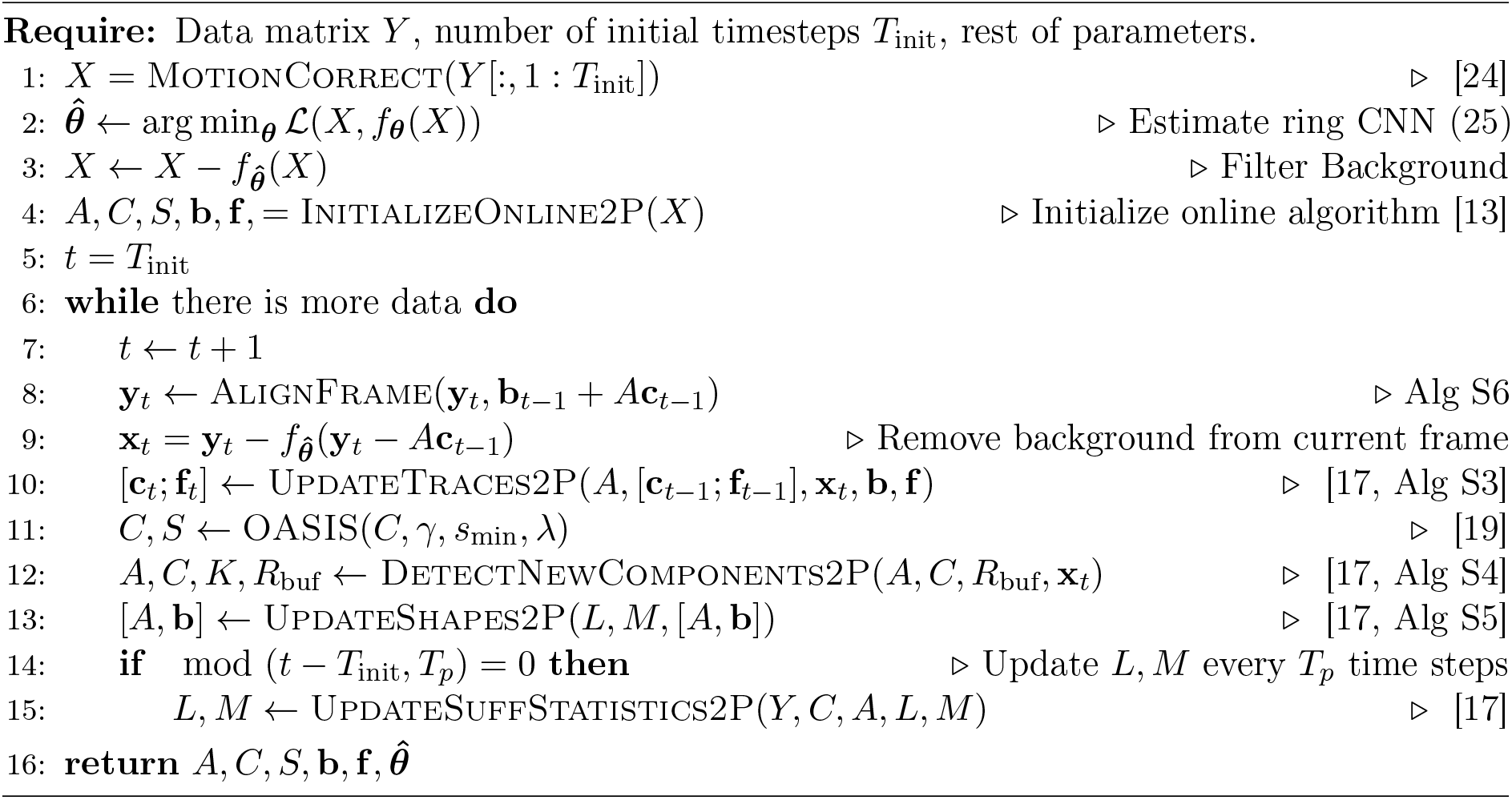
Online processing with a Ring-CNN background model

Once the background has been removed, online processing can be done using the standard online algorithm for two-photon data [17]. The process is summarized in Alg 2, where the suffix “2P” has been added to some routines to indicate their differences compared to the routines used in OnACID-E that are slightly more complicated due to their additional background treatment step. Note that although the focus of this paper is on online processing, the ring-CNN background model can also be used to derive an offline algorithm for microendoscopic 1p data.

### Online motion correction

Similarly to [17], online motion correction can be achieved by using the previously denoised frame **b**_*t*−1_ + *A***c**_*t*−1_ to derive a template for registering **y**_*t*_. In practice, we observed that this registration process is more robust to drift introduced by corrupt frames when an average of the past *N* denoised frames is used as a frame, with *N* ~ 50. As proposed in [13], passing both the template and the frame through a high pass spatial filter can suppress the strong background signal present in microendoscopic 1-photon data, and lead to more accurate computation of the alignment transformation. Rigid or piecewise rigid translations can be estimated as described in [24]. The inferred transformation is then applied to original frame **y**_*t*_. The process is summarized in Alg S6.

## Results

### Online analysis of 1p microendoscopic data using OnACID-E

We tested the online CNMF-E implementation of OnACID-E on in vivo microendosopic data from mouse dorsal striatum, with neurons expressing GCaMP6f. The data was acquired while the mouse was freely moving in an open field arena. The dataset consisted of 6000 frames at 10 Hz resolution (for further details refer to [5], for a second dataset see Fig S1). We initialized the online algorithm by running CNMF-E on the first 200 frames.

We illustrate OnACID-E in process in Fig 1. At the beginning of the experiment (Fig 1 left), only some components are active, as shown in panel A by the correlation image computed using the spatially filtered data [5], and most of these are detected by the algorithm (Fig 1B). As the experiment proceeds more neurons activate and are subsequently detected by OnACID-E (Fig 1 middle, right) which also tracks their activity across time (Fig 1C). See also Supplementary Video for further illustration.

**Fig 1.**
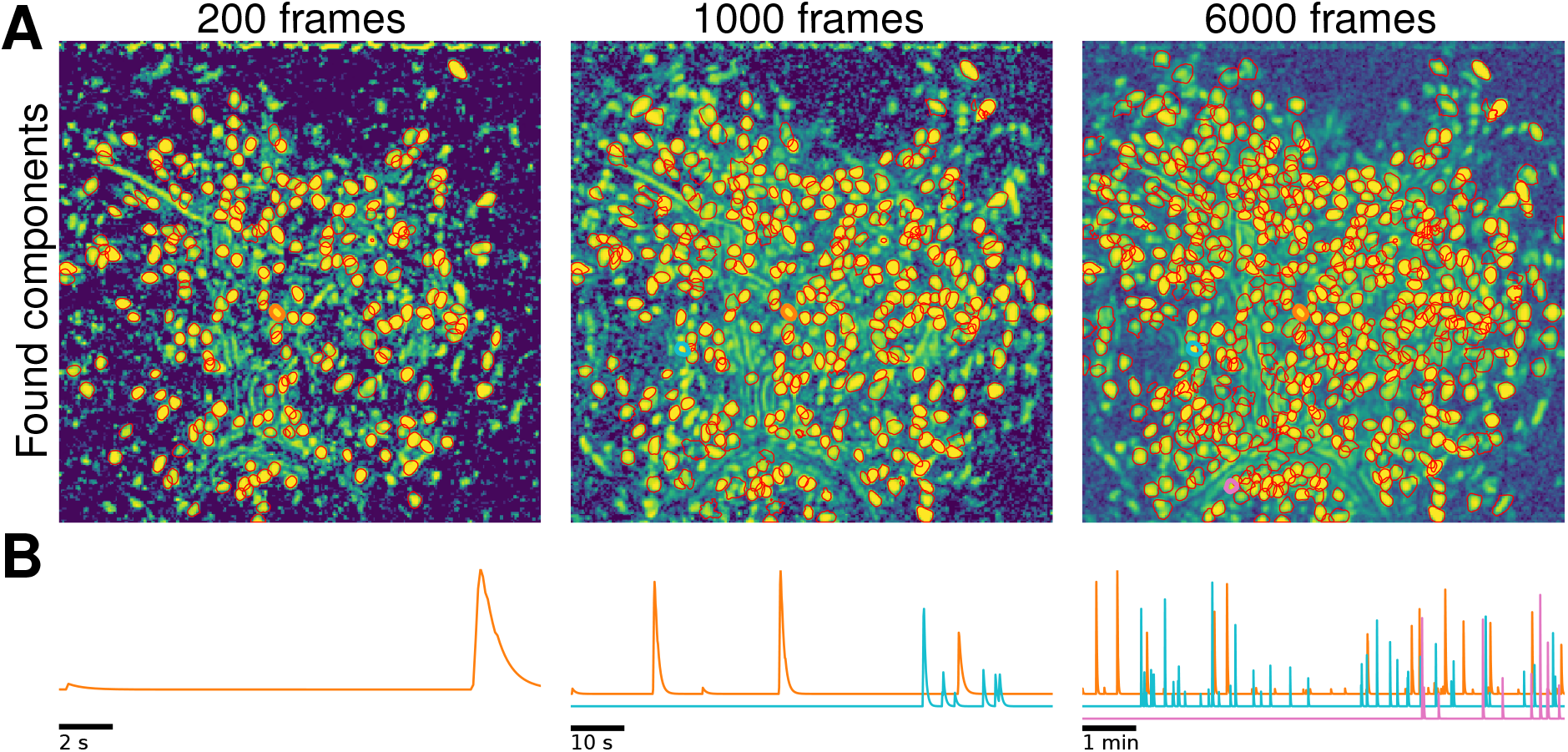
Illustration of the online data analysis process. Snapshots of the online analysis after processing 200 frames (left), 1000 frames (middle), and 6000 frames (right). **(A)** Contours of the components (neurons and processes) found by OnACID-E up to each snapshot point, overlaid over the local cross-correlation image of the spatially filtered data [5] at that point. **(B)** Examples of neuron activity traces (marked by corresponding colors in panel A). As the experiment proceeds, OnACID-E detects newly active neurons and tracks their activity.

### Comparison of OnACID-E with CNMF-E

In Fig 2 we report the results of the analysis using OnACID-E and compare to the results of CNMF-E with patches, i.e. the field of view (FOV) is split into smaller overlapping patches that are processed in parallel and combined at the end [13]. Both implementations detect similar components (Fig 2A) with an F1-score of 0.891 (0.875 if the variance summary image was used to detect new components). 506 components were found in common by both implementations. 48 and 76 additional components were detected by OnACID-E and CNMF-E respectively. Ten example temporal traces are plotted in Fig 2B. The first five are from neurons that have been detected in the initialization phase, the last five during online processing. While the neuron corresponding to the last trace was detected immediately once it became active, this wasn’t the case for the others (green traces). Low activity events can be too weak to trigger detection as new component, but are accurately captured once the existence of the neuron has already been established. Hence, performing a second online pass over the dataset recovers the entire activity traces (purple). The median correlation between the temporal traces of neurons detected by both implementations was 0.852.

**Fig 2.**
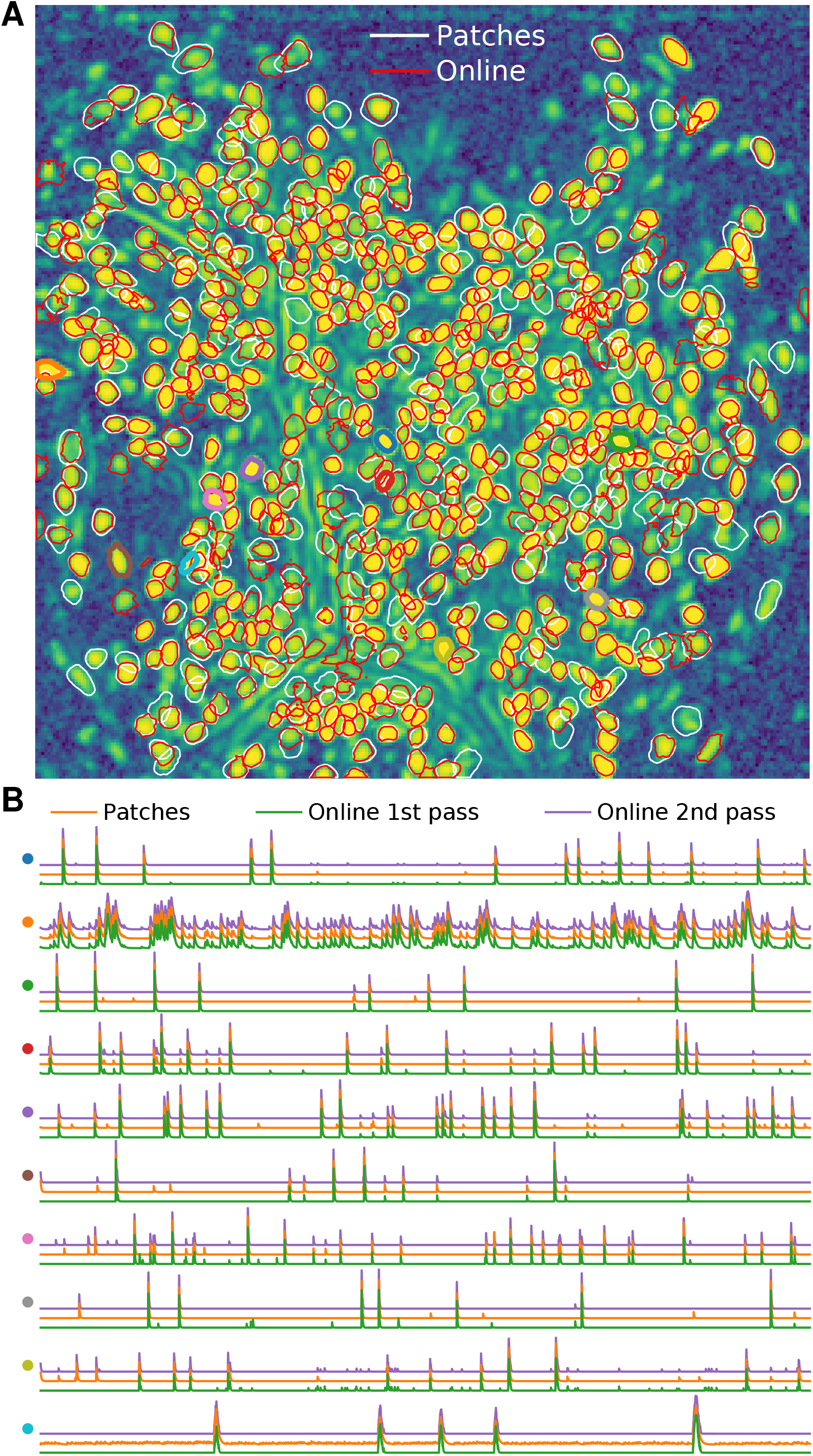
Comparison of OnACID-E with CNMF-E when analyzing microendoscopic 1-photon data. **(A)** Contour plots of all neurons detected by CNMF-E using patches (white) and OnACID-E (red), overlaid over the local cross-correlation image. Colors match the example traces shown in **(B)**, which illustrate the temporal components of ten example neurons detected by both implementations. The first five have been detected in the initialization phase, the last five during online processing.

We repeated the analysis on in-vivo microendosopic data from prefrontal cortex of a freely behaving mouse. This second dataset consisted of 9000 frames at 15 Hz resolution (for further details refer to [5]). Analogous results to Fig 2 are presented in Fig S1. The F1-score between components detected by OnACID-E and CNMF-E was 0.895.

We also performed the comparison on the simulated data from [5], in order to compare not only the offline and online method with each other but both with underlying ground truth, see Fig 3. Both implementations detect all components (Fig 2A) with a perfect F1-score of 1. We again show ten example temporal traces in Fig 3B. The overlaps 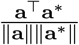 between true (**a**^∗^) and inferred (**a**) neural shape are reported in Fig 3C. While CNMF-E tends to capture the neural footprints more accurately, the inferred temporal components (that would be used in the subsequent analysis and are hence more important) are of similar quality, as the correlations with ground truth reveal (Fig 3D). The median correlation between the temporal traces of neurons detected by CNMF-E and ground truth was 0.996, for OnACID-E it was 0.993.

**Fig 3.**
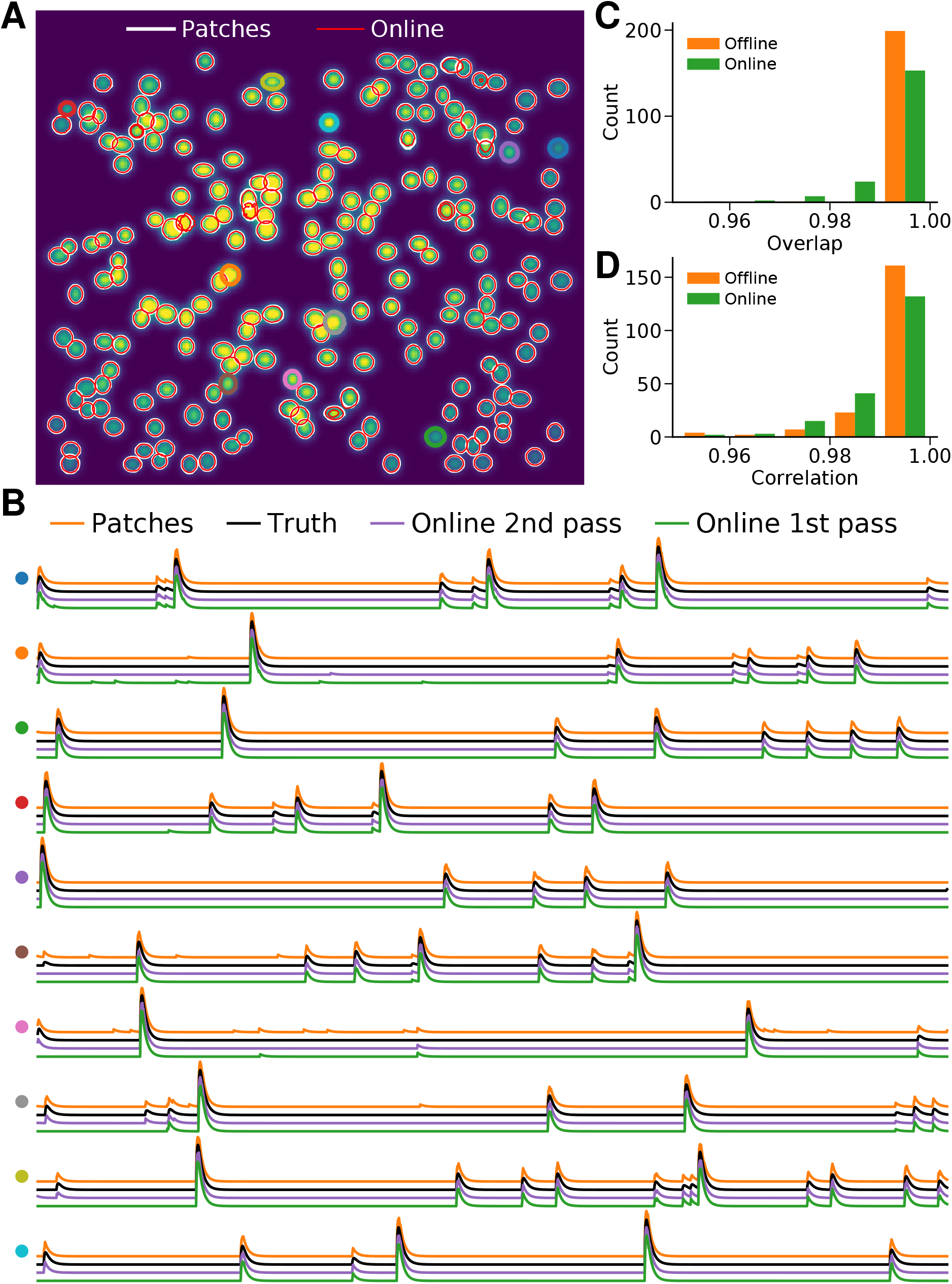
Comparison of OnACID-E with CNMF-E when analyzing simulated data. **(A)** Contour plots of all neurons detected by CNMF-E using patches (white) and OnACID-E (red), overlaid over the true neural shapes. Colors match the example traces shown in **(B)**, which illustrate the temporal components of ten example neurons detected by both implementations. The first five have been detected in the initialization phase, the last five during online processing. **(C)** Histogram of the overlaps between inferred and true neural shapes. **(D)** Histogram of the correlations between inferred and true neural fluorescence traces.

### Computational performance of OnACID-E

We examined the performance of OnACID-E in terms of processing time and memory requirements for the analyzed dataset presented above (Fig 4). The processing time discussed here excludes motion correction (which is highly efficient [24]), because the data was already motion corrected before hand. For the batch as well as the online algorithm we used the Python implementations provided by or added to CaImAn [13], respectively. The dataset was analyzed using a single node of a linux-based (CentOS) cluster with Intel Xeon Platinum 8168 CPU at 2.7 GHz (24 cores) and 768 GB of RAM. The same analysis was performed using merely a laptop (MacBook Pro 13”) with Intel Core i7-7567U CPU at 3.5 GHz (2 cores) and 16 GB of RAM.

The processing time of OnACID-E depends primarily on (i) the computational cost of tracking the temporal activity of discovered neurons, (ii) the cost of detecting and incorporating new neurons, and (iii) the cost of periodic updates of spatial footprints and background. Additionally, there is the one-time cost incurred for initialization. Fig 4A shows the cost of each of these steps for one epoch of processing. Initialization was performed by running CNMF-E on the first 200 frames, hence the sudden jump at 200 processed frames in Fig 4A. The cost of detecting and incorporating new components remains approximately constant across time and is dependent on the number of candidate components at each time step. In this example three candidate components were used per frame. As noted in [13], a higher number of candidate components can lead to higher recall in shorter datasets at a moderate additional computational cost.

The cost of tracking components can be kept low due to simultaneous vectorized updates, and increases only mildly over time as more components are found by the algorithm, cf. Fig 4B. Finally, it is particularly noteworthy that the total processing time was smaller than the duration of the recording. Processing times are slightly smaller when the variance summary image is used (Fig S1), but at the expense of the reported smaller, albeit similar, F1 score.

Fig 4C shows the memory usage as function of processing time and compares to CNMF-E with or without splitting the FOV into patches. Sixteen patches of size 96×96 were used and processed simultaneously in parallel. OnACID-E does not only use less memory but is even faster than CNMF-E without patches. Processing can be faster using patches, however, this gain comes at the cost of enormous memory requirements and necessitates a powerful computing environment, such as for example the cluster node we used. These requirement can be mitigated at the expense of longer processing times by processing not all patches in parallel, as the additional orange markers in Fig 4C for 8, 6, 4, 3, 2 and 1 parallel processes show.

When the analysis was performed on a laptop (with four threads) not all, but merely up to four of the total sixteen patches, could be processed simultaneously in parallel. Fig 4C shows that whereas processing in patches was marginally faster and less memory consuming than processing the entire FOV, both are clearly outperformed with regard to computing time and memory requirements by OnACID-E. It required less memory than the size of the whole data, here 1.5 GB (for single-precision float), and about an order of magnitude less memory than CNMF-E. These results would be even more pronounced for longer datasets because the memory consumption remains nearly constant as time progresses and is thus independent of the number of recorded frames. Online processing on the laptop took about the same time as on the cluster node. OnACID-E strikes the best balance between memory consumption and processing time, making it in particular suitable for processing of long datasets without the need for high performance hardware.

### Performance of the Ring CNN approach

For comparison purposes we also tried the online analysis of the same dorsal striatum dataset, using the Ring-CNN background model (Eq (22)) with two kernels of width 5 pixels. The model was trained on the first 500 frames (400 frames for training and 100 for validation) using a quantile loss function (Eq (26)) with *q* = 0.02, using stochastic gradient descent with the ADAM optimizer. After initialization every frame was passed through the learned model to remove its background and was subsequently processed using the CaImAn Online algorithm [13] with a rank-2 background. During this phase, the background model was kept constant with no additional training, which resulted in faster processing. This was possible because the background model had already converged to a stable value during initialization because of the smaller number of parameters needed to be learned due to the large level of weight sharing. Moreover, the increased width of the filter increased the statistical power of the model making it less sensitive to outliers, and thus aiding faster convergence. Three epochs were used to process the dataset, with the third epoch being used only to track the activity of the existing neurons (and not to detect new components). After the online processing was done, the identified components were merged, and then screened for false positive using the tests employed in the CaImAn package [13].

Lacking a “ground truth” benchmark we compared its performance against the CNMF-E algorithm [5]. The results of the analysis are summarized in Fig 5. The algorithms displayed a high level of agreement (black contours in Fig 5A) with F1-score 0.848 (precision 0.81 and recall 0.888 treating the CNMF-E predictions as “ground truth”). While the agreement between the ring CNN appoach and CNMF-E was lower compared to the agreement between OnACID-E and CNMF-E, this cannot be readily interpreted as underperformance of the ring CNN approach. For example, the ring CNN approach identified several components that have a clear spatial footprint in the correlation image of the spatially filtered data (some examples are highlighted by the orange arrows).

**Fig 5.**
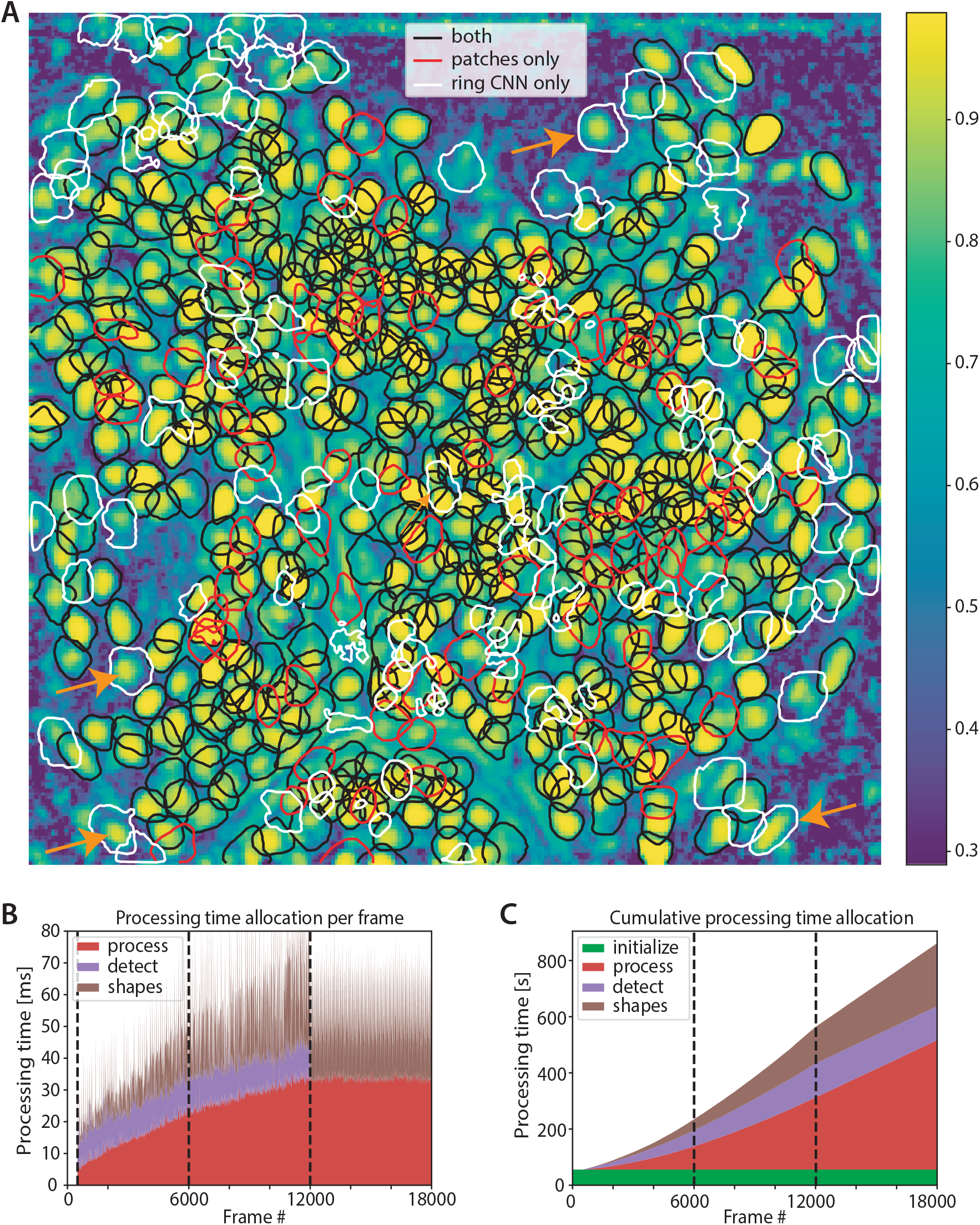
Performance of online approach using a ring CNN background model. **(A)** Contour plots of all neurons detected by the ring CNN approach and CNMF-E using patches overlaid over the local cross-correlation image. The two approaches have a high level of similarity (black contours, F1-score 0.845), with several components identified only by one algorithm (red contours, CNMF-E only, white contours ring, CNN only). At least some of the contours identified only by the ring CNN model appear to correspond to actual neurons (orange arrows). Processing speed per frame **(B)** and cumulatively **(C)** for the ring CNN approach. By reducing the background extraction to a simple, GPU-implementable, filtering operation and estimating it only during initialization, the ring CNN approach can achieve high processing speeds for every frame **(B)**, and run a complete epoch on the data faster than OnACID-E **(C)**. Moreover, it can distribute the computational load evenly amongst all frames making it useful for real time applications.

The computational performance of the ring CNN approach is shown in Fig 5B-C. In addition to a computing cluster node, an NVIDIA Tesla V100 SXM2 32GB GPU was deployed to estimate the background model and subsequently apply it. Overall initialization on the 500 frames required around 53s, roughly equally split between estimating and applying the background model, and performing “bare initialization” [13] on the background extracted to find 50 components and initialize the rank-2 background. After that processing was very fast for every frame (Fig 5B) with no computational bottlenecks (as opposed to OnACID-E where updating the background can take significant resources). Overall, the first epoch of processing was completed in 210s (Fig 5C), a factor of 2 improvement over OnACID-E (even without background updating for OnACID-E). A closer comparison between Fig 4A and Fig 5C indicates that the ring CNN approach is faster than OnACID-E for detecting new components, but slower during “tracking”. The reason for that, is that “tracking” in the ring CNN approach includes the background removing step (which requires data transfer to and from the GPU). However, once this step is done, no additional background treatment is required, which speeds up the detection step significantly. More importantly, this allowed a distributed update of shapes amongst all frames (Fig 5B) which kept the processing speed for *each* frame above the acquisition rate of 10Hz, thus achieving real time processing. Since the initialization step can be performed in mini-batches the GPU memory requirements remain limited. After that, online processing is deployed on a frame by frame basis which keep the memory requirements at similar levels compared to OnACID-E (data not shown).

## Discussion

We presented an online method to process 1-photon microendoscopic video data. Our modeling assumptions are the same as in the popular offline method CNMF-E; however, our online formulation yields a more efficient yet similarly accurate method for the extraction of in vivo calcium signals. A major bottleneck for processing microendoscopic data has been the amount of memory required by CNMF-E. Our online approach solves this issue since it reduces the memory footprint from scaling linearly with the duration of the recording to being constant. We also provided an additional variant that uses a convolutional based background model that aims to exploit the location invariant properties of the point spread function. This approach enables the estimation of a stable background model by using just an initial portion of the data. As a result, it can lead to faster processing and also be coupled to 2-photon processing algorithms by using this model to remove the background from each frame as a preprocessing step.

For detecting centroids of new sources OnACID-E examines a static image. Following [17], such an image can be obtained by computing the variance across time of the spatially smoothed residual buffer. As an additional option to obtain a static summary image we added the computation of peak-to-noise ratio and local cross correlation across time of the residual buffer, following the proposal of [5]. For efficiency, this computation is performed online using incremental updates. While both work very well in practice, different approaches for detecting neurons in static images or in a short residual buffer could potentially be employed here, e.g. dictionary learning [25], combinatorial clustering [26] or deep neural networks [27, 28]. However, these approaches likely come with higher computational cost, and – having been developed for offline processing – would probably need to be modified for data streams, and in the case of neural networks be retrained.

Similarly to [17], our current implementation screens the candidate components for quality using some quantitative measures and thresholds. For 2-photon data [13] suggested to use a neural net classifier instead for better accuracy. Training a neural network requires labelled data, which is currently not publicly available for 1-photon microendoscopic video data. Once labelled ground truth data is available, a neural network could be trained on it and OnACID-E be readily augmented to use this classifier. Such ground truth data would also enable to thoroughly benchmark different source extraction algorithms and their implementations.

Apart from enabling rapid and memory efficient analysis of microendoscopic 1-photon data, our online pipeline also facilitates closed-loop behavioral experiments that analyze data on-the-fly to guide the next experimental steps or to control feedback. The current implementation of OnACID-E is already faster than real time on average. On a per-frame basis the processing speed exceeds the data rate for the majority of frames, and only when the periodic updates of sufficient statistics, shapes, and background are performed can the speed drop below the data rate. This can be ameliorated by using a larger initialization batch for OnACID-E. Once enough initial data has been seen and processed, the computationally expensive search for components as well as the spatial footprint and background updates can be turned off, because all regions of interest have been detected and their shapes as well as the background converged to stable values. Further, as presented, this compromise can be avoided altogether by endowing the background with a convolutional structure that enables faster convergence in the background estimation. This subsequently enables updating of spatial footprints in a distributed sense, while maintaining faster than real time processing rates at *every* frame by keeping the ability to detect and incorporate new components.

## Availability

We provide a Python implementation of our algorithm online within CaImAn, an open-source library for calcium imaging data analysis (https://github.com/flatironinstitute/CaImAn) [13]. Our work extends the library to enable online processing of microendoscopic 1-photon data.

## Acknowledgments

We would like to thank Kris Pan for useful discussions.

## Supplementary Material

Here we present in pseudocode the various steps of the online processing pipeline. For ease of exposition, some details and speedup tricks used in the actual implementation have been omitted, such as the online update of the summary image used for neuron detection or the spatial decimation of the background.

**Algorithm S1.**
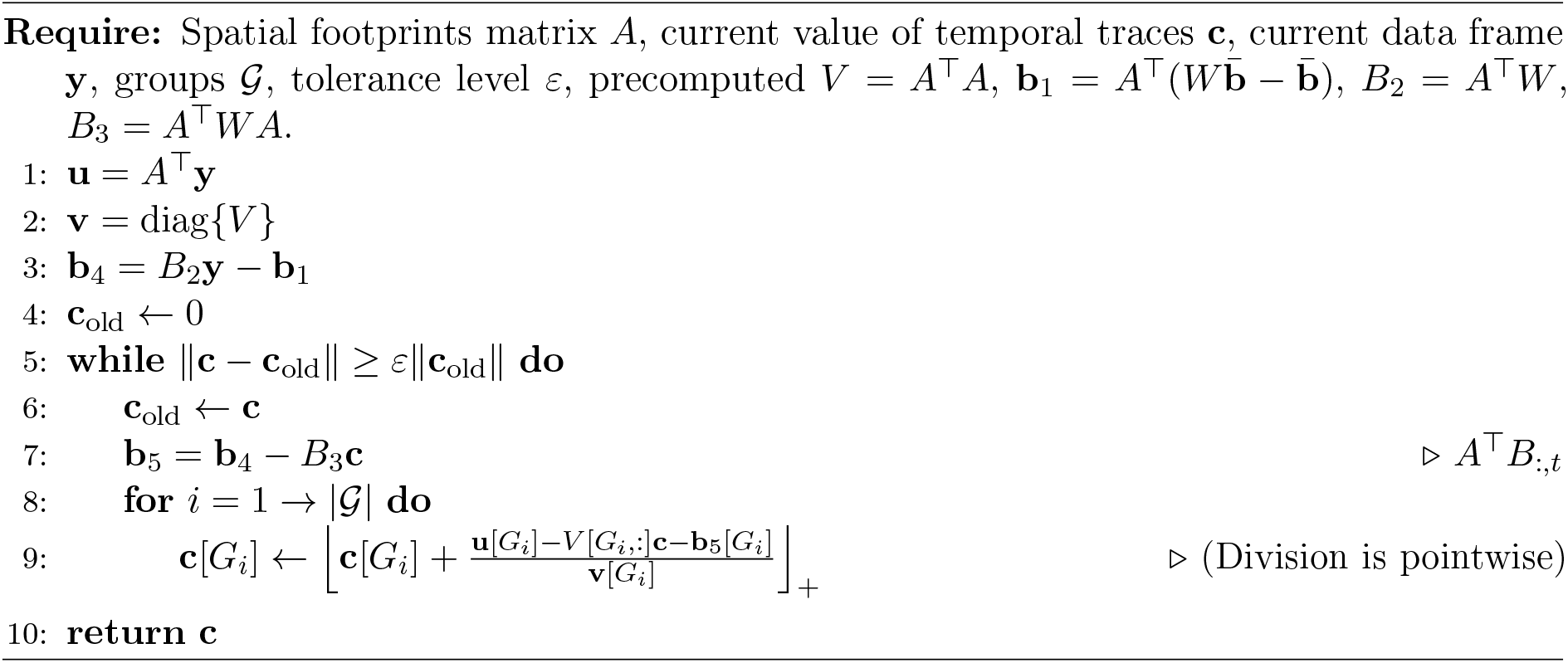
UpdateTraces

**Algorithm S2.**
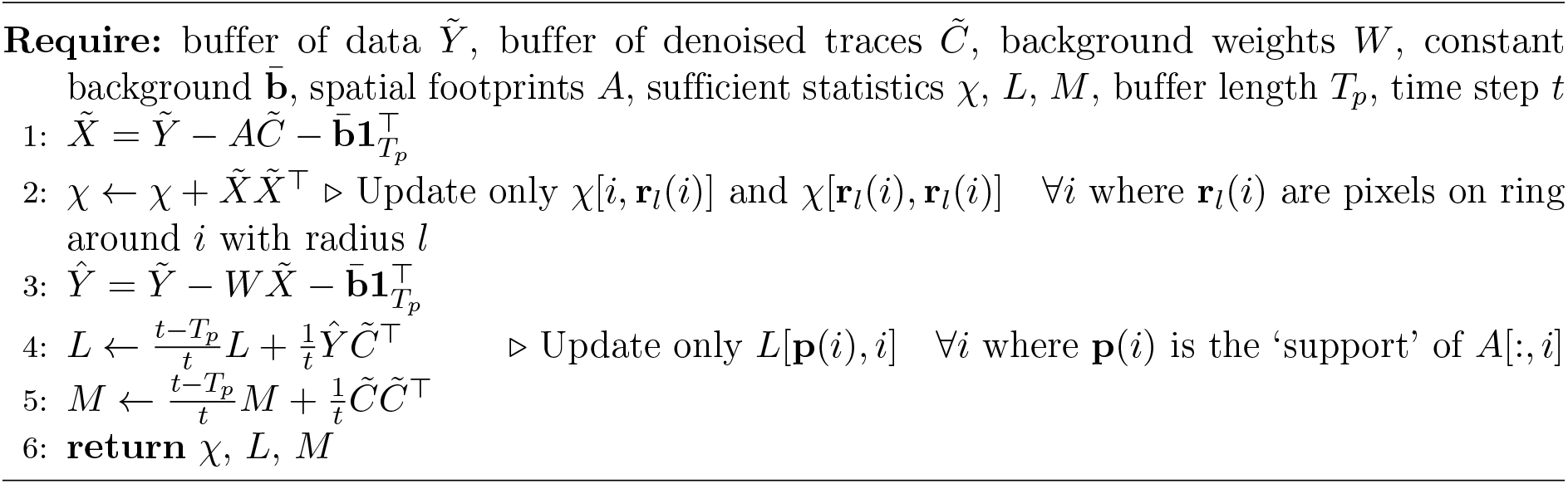
UpdateSuffStatisticsl

**Algorithm S3.**
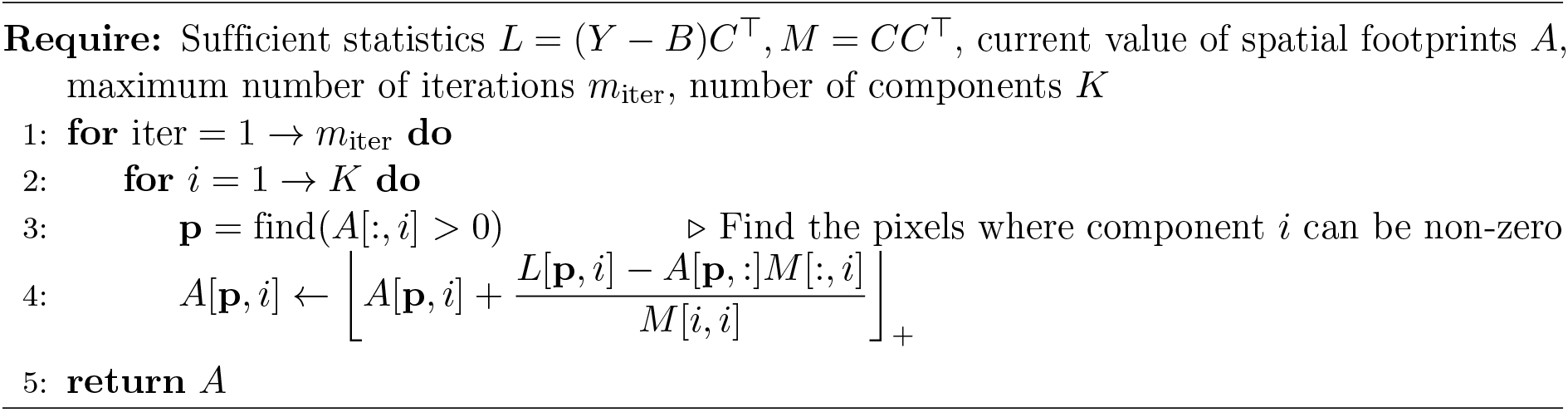
UpdateShapes

**Algorithm S4.**
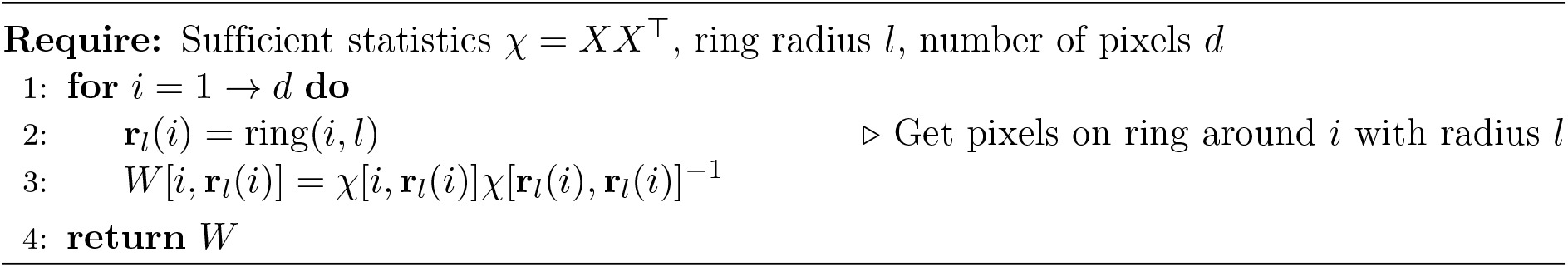
UpdateBackground

**Algorithm S5.**
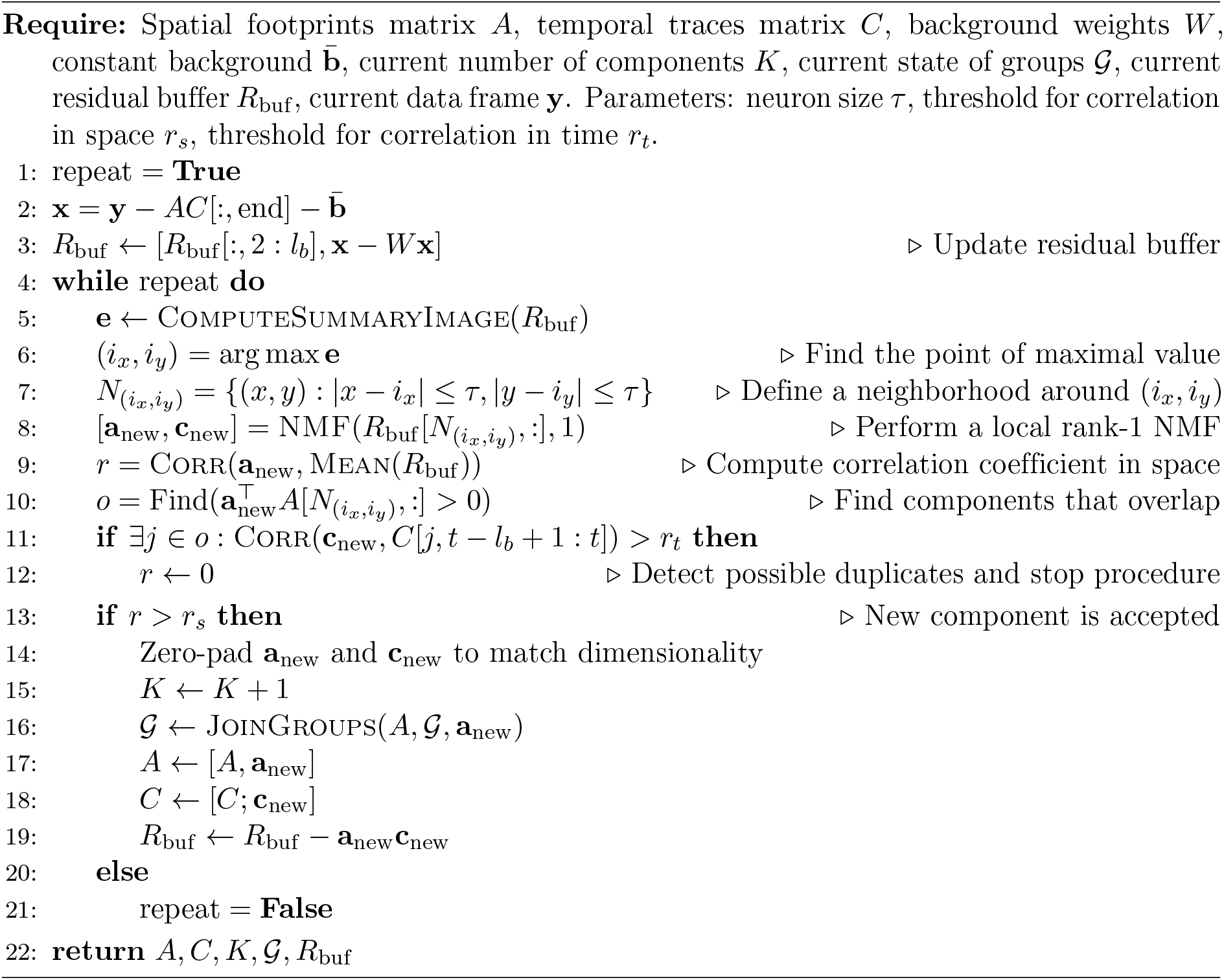
DetectNewComponents

**Algorithm S6.**
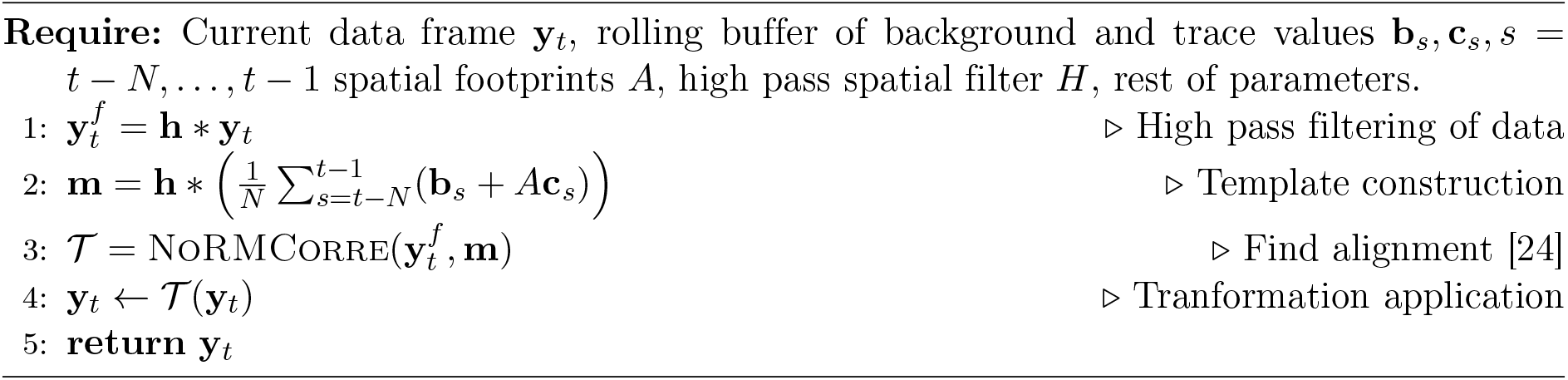
AlignFrame

**Fig S1.**
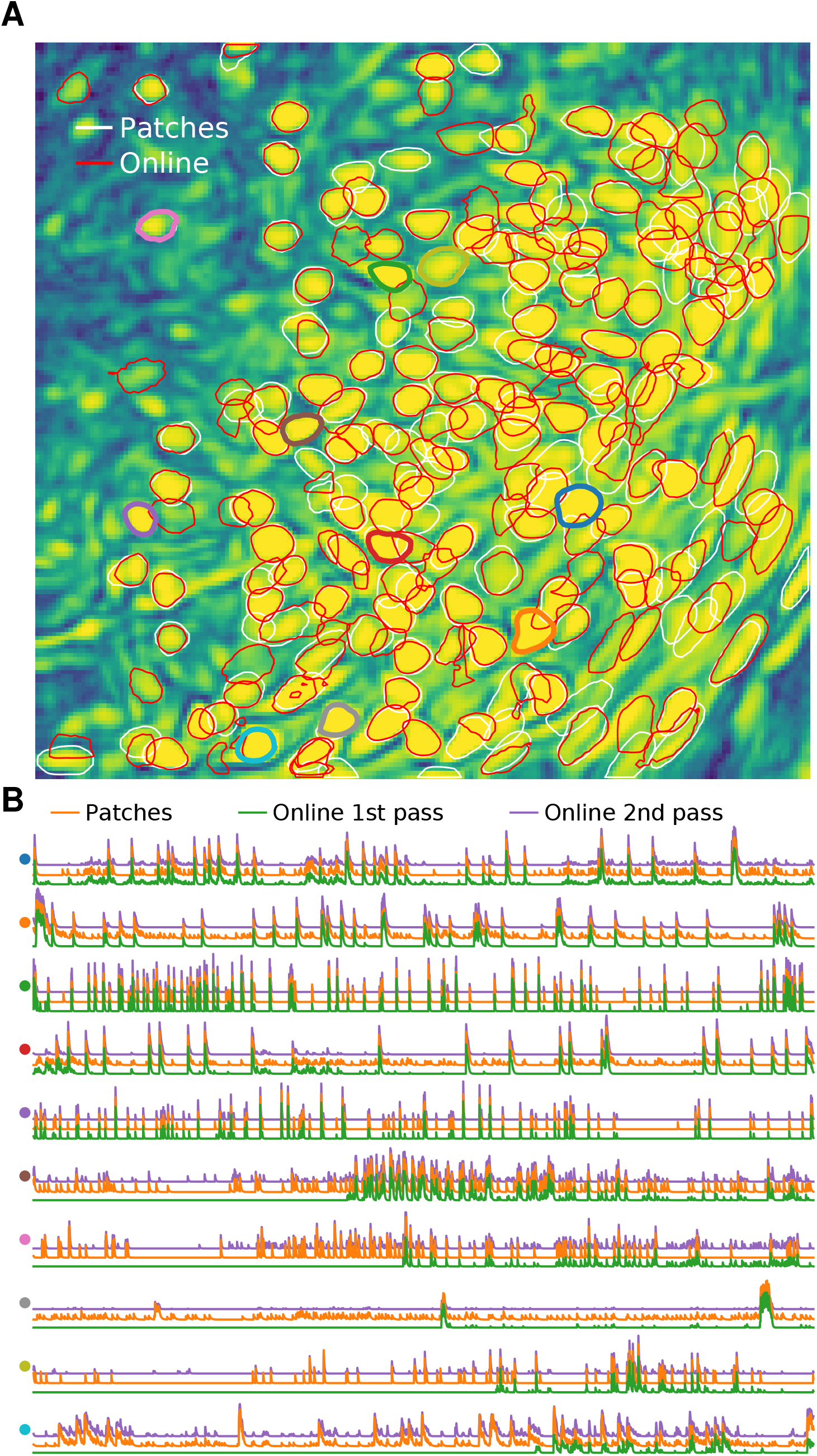
Comparison of OnACID-E with CNMF-E on data from prefrontal cortex. Analogous plots to Fig 2 for in-vivo microendosopic data from prefrontal cortex of a freely behaving mouse.

**Fig S2.**
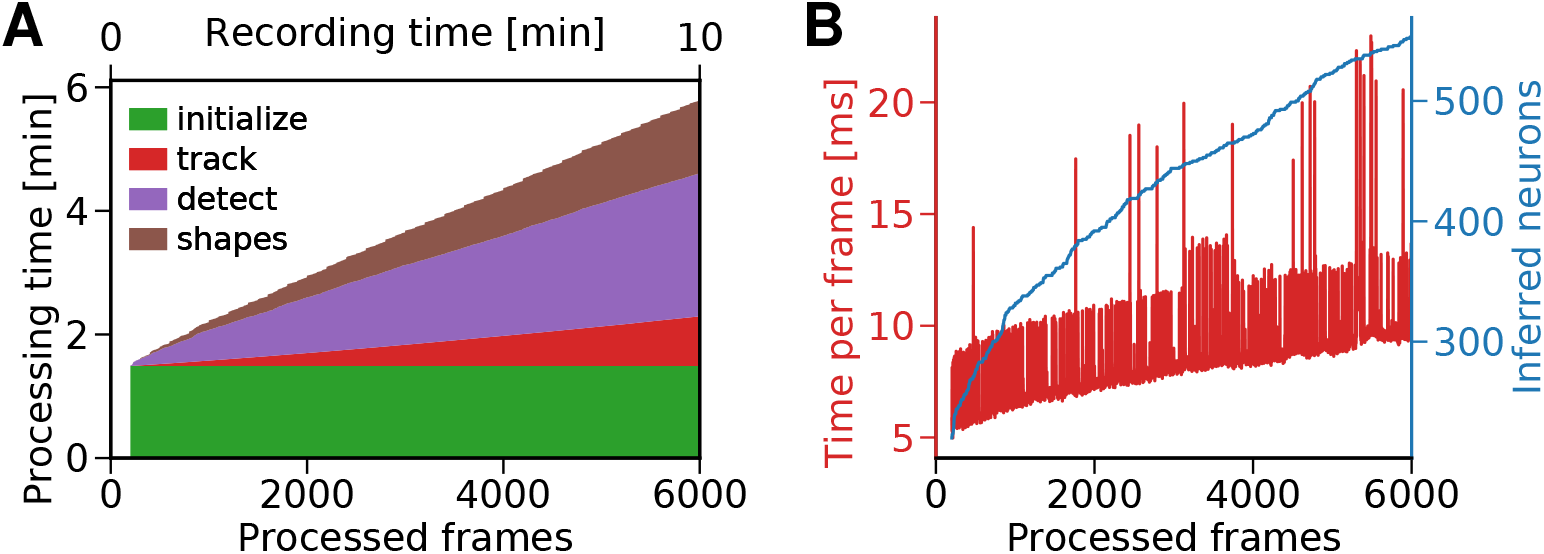
Processing time of OnACID-E using the variance summary image. Analogous plots to Fig 4A and B.

## Supplementary Video

### Depiction of OnACID-E

Top left: Raw data. Top right: Inferred activity (without background). Bottom left: Corr*PNR summary image (see Methods) and accepted regions for new components (magenta squares). Bottom right: Reconstructed activity.

